# Epidemic establishment and cryptic transmission of Zika virus in Brazil and the Americas

**DOI:** 10.1101/105171

**Authors:** N. R. Faria, J. Quick, I. Morales, J. Thézé, J.G. Jesus, M. Giovanetti, M. U. G. Kraemer, S. C. Hill, A. Black, A. C. da Costa, L.C. Franco, S. P. Silva, C.-H. Wu, J. Raghwani, S. Cauchemez, L. du Plessis, M. P. Verotti, W. K. de Oliveira, E. H. Carmo, G. E. Coelho, A. C. F. S. Santelli, L. C. Vinhal, C. M. Henriques, J. T. Simpson, M. Loose, K. G. Andersen, N. D. Grubaugh, S. Somasekar, C. Y. Chiu, J. E. Muñoz-Medina, C. R. Gonzalez-Bonilla, C. F. Arias, L. L. Lewis-Ximenez, S.A. Baylis, A. O. Chieppe, S. F. Aguiar, C. A. Fernandes, P. S. Lemos, B. L. S. Nascimento, H. A. O. Monteiro, I. C. Siqueira, M. G. de Queiroz, T. R. de Souza, J. F. Bezerra, M. R. Lemos, G. F. Pereira, D. Loudal, L. C. Moura, R. Dhalia, R. F. França, T. Magalhães, E. T. Marques, T. Jaenisch, G. L. Wallau, M. C. de Lima, V. Nascimento, E. M. de Cerqueira, M. M. de Lima, D. L. Mascarenhas, J. P. Moura Neto, A. S. Levin, T. R. Tozetto-Mendoza, S. N. Fonseca, M. C. Mendes-Correa, F.P. Milagres, A. Segurado, E. C. Holmes, A. Rambaut, T. Bedford, M. R. T. Nunes, E. C. Sabino, L. C. J. Alcantara, N. Loman, O. G. Pybus

## Abstract

Zika virus (ZIKV) transmission in the Americas was first confirmed in May 2015 in Northeast Brazil^1^. Brazil has the highest number of reported ZIKV cases worldwide (>200,000 by 24 Dec 2016^2^) as well as the greatest number of cases associated with microcephaly and other birth defects (2,366 confirmed cases by 31 Dec 2016^2^). Following the initial detection of ZIKV in Brazil, 47 countries and territories in the Americas have reported local ZIKV transmission, with 24 of these reporting ZIKV-associated severe disease^3^. Yet the origin and epidemic history of ZIKV in Brazil and the Americas remain poorly understood, despite the value of such information for interpreting past and future trends in reported microcephaly. To address this we generated 54 complete or partial ZIKV genomes, mostly from Brazil, and report data generated by the ZiBRA project – a mobile genomics lab that travelled across Northeast (NE) Brazil in 2016. One sequence represents the earliest confirmed ZIKV infection in Brazil. Joint analyses of viral genomes with ecological and epidemiological data estimate that ZIKV epidemic was present in NE Brazil by March 2014 and likely disseminated from there, both nationally and internationally, before the first detection of ZIKV in the Americas. Estimated dates of the international spread of ZIKV from Brazil indicate the duration of pre-detection cryptic transmission in recipient regions. NE Brazil’s role in the establishment of ZIKV in the Americas is further supported by geographic analysis of ZIKV transmission potential and by estimates of the virus’ basic reproduction number.

**One Sentence Summary:** Virus genomes reveal the establishment of Zika virus in Brazil and the Americas, and provide an appropriate timeframe for baseline (pre-Zika) microcephaly in different regions.

---

Previous phylogenetic analyses indicated that the ZIKV epidemic was caused by the introduction of a single Asian genotype lineage into the Americas around late 2013, at least one year before its detection there^4^. An estimated 100 million people in the Americas are predicted to be at risk of acquiring ZIKV once the epidemic has reached its full extent^5^. However, little is known about the genetic diversity and transmission history of the virus in different regions in Brazil^6^. Reconstructing ZIKV spread from case reports alone is challenging because symptoms (typically fever, headache, joint pain, rashes, and conjunctivitis) overlap with those caused by co-circulating arthropod-borne viruses^7^ and due to a lack of nationwide ZIKV-specific surveillance in Brazil before 2016.

**Fig. 1.**
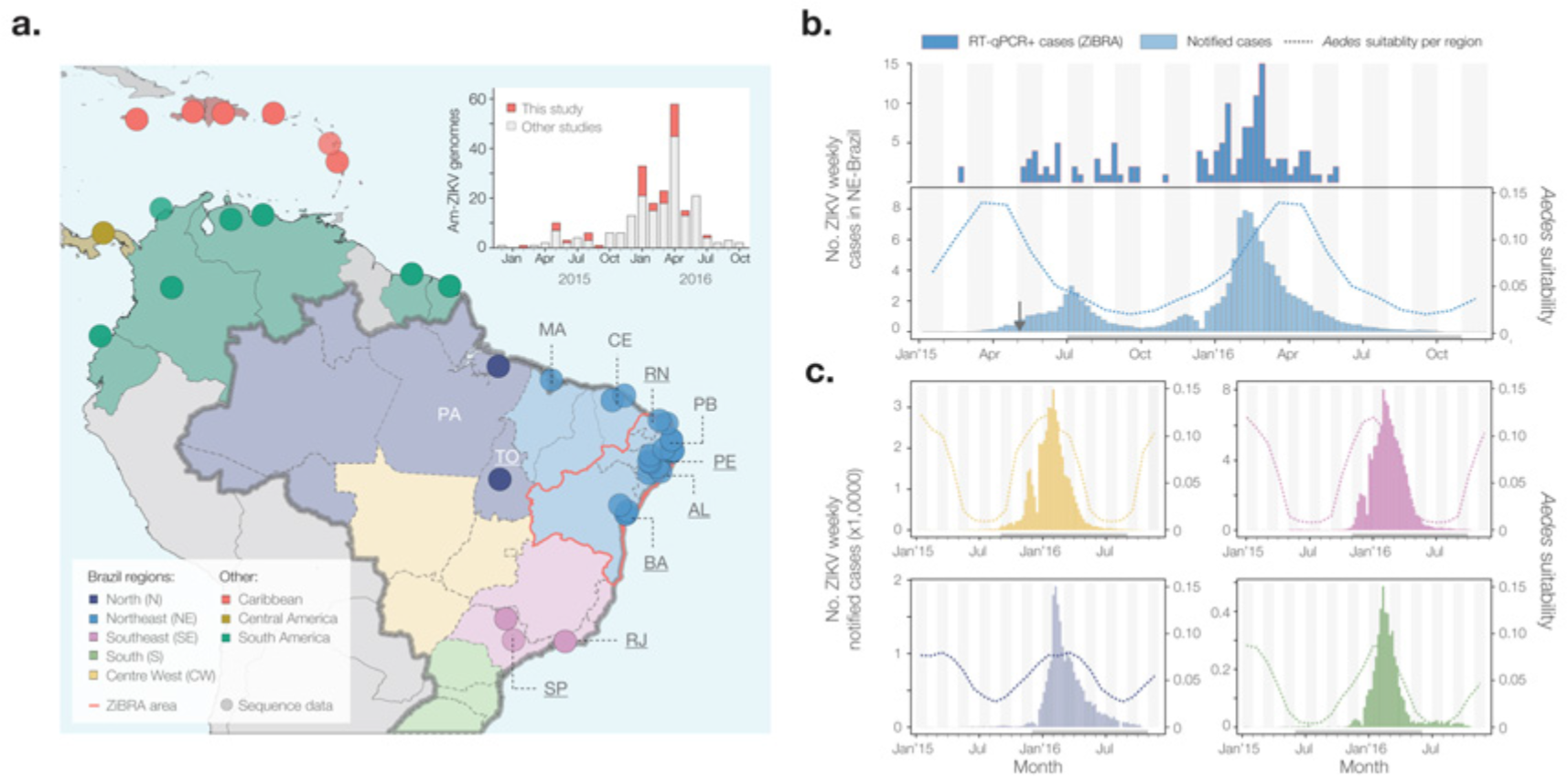
Geographic and temporal distribution of ZIKV in Brazil. **a.** Location of sampling of genome sequences from Brazil and in the Americas. Federal states in Brazil are coloured according to 5 geographic regions (see bottom left inset). A red contour line surrounds the federal states surveyed during June 2016 by the ZiBRA mobile lab. Two letter state codes are as follows: PA: Pará, MA: Maranhão, CE: Ceará, TO: Tocantins, RN: Rio Grande do Norte, PB: Paraíba, PE: Pernambuco, AL: Alagoas, BA: Bahia, RJ: Rio de Janeiro, SP: São Paulo. Underlined states represent those from which sequences in this study were generated. Non-underlined states represent those from which publicly available sequences were collated. **b.** Confirmed and notified ZIKV cases in NE Brazil. The upper panel shows the temporal distribution of RT-qPCR+ cases (n=181) detected during the ZiBRA journey. Only confirmed cases for which the exact collection date was known (138 out of 181) are included. The lower panel shows notified ZIKV cases in NE Brazil between 01 Jan 2015 and 19 Nov 2016 (n = 122,779). The dashed line represents the average climatic vector suitability score for NE Brazil (see Methods). The vertical arrow indicates date of ZIKV confirmation in the NE Brazil/Americas ^1^. **c.** Notified ZIKV cases in the Centre-West, Southeast, North and South regions of Brazil (clockwise from top left). As before, the dashed lines represent the average climatic vector suitability score for each region. The R^2^, P-value and estimated lag (T) values of the model used to compare notified cases with vector suitability scores are provided in Extended Data Table 2. Grey horizontal bars below each time series indicate the time period for which the correlation between suitability and ZIKV notified cases was highest.

To address this we undertook a collaborative investigation of ZIKV molecular epidemiology in Brazil, including results from a mobile genomics laboratory that travelled through NE Brazil during June 2016 (the ZiBRA project; http://www.zibraproject.org). Of five regions of Brazil (Fig. 1a), the Northeast region (NE Brazil) has the most notified ZIKV cases (40% of Brazilian cases) and the most confirmed microcephaly cases (76% of Brazilian cases, to 31 Dec 2016^2^), raising questions about why the region has been so severely affected^8^. Further, NE Brazil is the most populous region of Brazil with the potential for year-round ZIKV transmission^9^. With the support of the Brazilian Ministry of Health and other institutions (**Acknowledgements**), the ZiBRA lab screened 1330 samples (almost exclusively serum or blood) from patients residing in 82 municipalities across five federal states in NE Brazil (Fig. 1; Extended Data Table 1). Samples provided by the central public health laboratory of each state (LACEN) and FIOCRUZ were screened for the presence of ZIKV by real time quantitative PCR (RT-qPCR).

On average, ZIKV viremia persists for 10 days after infection; symptoms develop ∼6 days after infection and can last 1-2 weeks^10^. In line with previous observations in Colombia^11^, we found that the RT-qPCR+ samples in NE Brazil were, on average, collected only two days after onset of symptoms. The median RT-qPCR cycle threshold (Ct) value of positive samples was correspondingly high, at 36 (Extended Data Fig. 1). For NE Brazil, the time series of RT-qPCR+ cases was positively correlated with the number of weekly-notified cases (Pearson’s ρ=0.62; Fig. 1b).

The ability of the mosquito vector *Aedes aegypti* to transmit ZIKV is determined by ecological factors that affect adult survival, viral replication, and infective periods^12^. To investigate the receptivity of each Brazilian region to ZIKV transmission, we used a measure of vector climatic suitability derived from monthly temperature, relative humidity, and precipitation data^9^. Using linear regression we find that, for each Brazilian region, there is a strong association between estimated climatic suitability and weekly notified cases (Figs. 1b,1c; adjusted R^2^>0.84, P<0.001; Extended Data Table 2). Similar to previous findings obtained for dengue virus outbreaks^13,14^, notified ZIKV cases lag climatic suitability by ∼4 to 6 weeks in all regions, except NE Brazil, where no time lag is evident. Despite these associations, numbers of notified cases should be interpreted cautiously because (i) co-circulating dengue and Chikungunya viruses exhibit symptoms similar to ZIKV, and (ii) the Brazilian case reporting system has evolved through time (see **Methods**). We estimated the basic reproductive numbers (R_0_) for ZIKV in each Brazilian region from the weekly notified case data and found that R_0_ is high in NE Brazil (R_0_∼3 for both epidemic seasons; Extended Data Table 3). Although our R_0_ values are approximate, in part due to spatial variation in transmission across the large regions analysed here, they are consistent with previous estimates from a variety of approaches^15,16^.

**Fig. 2.**
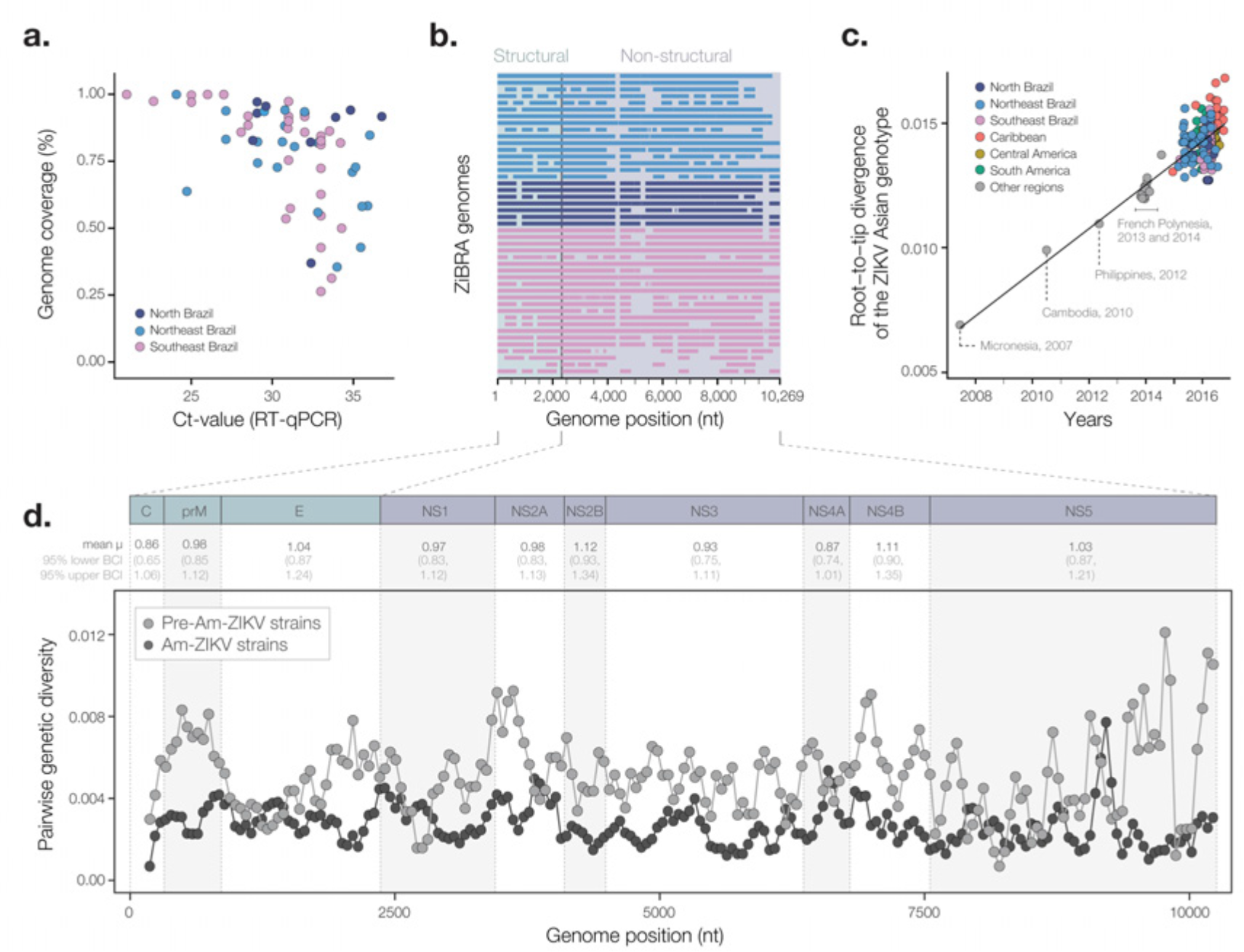
Zika virus genetic diversity and sequencing statistics. **a.** Plot showing the percentage of ZIKV genome sequenced against RT-qPCR Ct-value for each sample. Each circle represents a sequence sample recovered from an infected individual in Brazil and is colour coded according to the location of sampling. **b**. Illustration of sequencing coverage across the ZIKV genome for the ZiBRA sequences, including data generated by both the mobile laboratory and by static labs. **c.** Regression of sequence sampling dates against root-to-tip genetic distances in ML phylogenetic tree of the Asian-ZIKV lineage. This plot excludes P6-740 (the oldest Asian-ZIKV strain, collected in Malaysia in 1966). A comparable analysis that includes P6-740 is shown in Extended Data Fig. 2. d. Average pairwise genetic diversity of the PreAm-ZIKV strains (grey line) and of the Am-ZIKV lineage (black line), calculated using a sliding window of 300 nucleotides with a step size of 50 nucleotides. Above the plot, estimated per-gene rates of evolution are shown (mean = µ and 95% Bayesian credible intervals = BCIs) are shown in units of 10^−3^ substitutions per site per year.

Encouraged by the utility of portable genomic technologies during the West African Ebola virus epidemic^17^ we used our openly-developed protocol^18^ to sequence ZIKV genomes directly from clinical material using MinION DNA sequencers. We were able to generate virus sequences within 48 hours of the mobile lab’s arrival at each LACEN. In pilot experiments using a cultured ZIKV reference strain^19^ our protocol recovered 98% of the virus genome (Extended Data Fig. 2). However, due to low viral copy numbers in clinical samples (Extended Data Fig. 1) many sequences exhibited incomplete genome coverage and were subjected to additional sequencing efforts in static labs once fieldwork was completed. Whilst average genome coverage was higher for samples with lower Ct-values (85% for Ct<33; Fig. 2a, Extended Data Table 4), samples with higher Ct values had highly variable coverage (mean=72% for Ct≥33 (Fig. 2a). Unsequenced genome regions were non-randomly distributed (Fig. 2b), suggesting that the efficiency of PCR amplification varied among primer pair combinations. We generated 36 near-complete or partial genomes from the NE, SE and N regions of Brazil, supplemented by 9 sequences from samples from Rio de Janeiro municipality. To further elucidate Zika virus transmission in the Americas, we include in our analyses 5 new ZIKV complete genomes from Colombia and 4 from Mexico. Further, we append to our dataset 115 publicly available sequences as well as 85 additional genomes from a companion paper ^20^. The dataset used for genetic analysis thus comprised 254 complete and near complete genomes; 241 of which were sampled in the Americas (**Methods**).

The American ZIKV epidemic comprises a single founder lineage^4,21,22^ (hereafter termed Am-ZIKV) derived from Asian genotype viruses (hereafter termed PreAm-ZIKV) from Southeast Asia and the Pacific^4^. A sliding window analysis of pairwise genetic diversity along the ZIKV genome shows that the diversity of PreAm-ZIKV strains is on average ∼2.1-fold greater than Am-ZIKV viruses (Fig. 2d), reflecting a longer period of ZIKV circulation in Asia and the Pacific than in the Americas. Genetic diversity of the Am-ZIKV lineage will increase in future and updated diagnostic assays are recommended to guarantee RT-qPCR sensitivity^23^.

It has been suggested that recent ZIKV epidemics may be causally linked to a higher apparent evolutionary rate for the Asian genotype than the African genotype^24,25^. However, such comparisons are confounded by an inverse relationship between the timescale of observation and estimated viral evolutionary rates^26^. Regression of sequence sampling dates against root-to-tip genetic distances indicates that molecular clock models can be applied reliably to the Asian-ZIKV lineage (Fig. 2c; Extended Data Figs. 3 and 4). We estimate the whole genome evolutionary rate of Asian ZIKV to be 0.97×10^−3^ substitutions per site per year (s/s/y; 95% Bayesian credible interval, BCI=0.87−1.01×10^−3^), consistent with other estimates for the Asian genotype^4,25^. We found no significant differences in evolutionary rates among ZIKV genome regions (Fig. 2d). The estimated d_N_/d_S_ ratio of the Am-ZIKV lineage is low (0.11, 95% CI= 0.10-0.13), as observed for other vector-borne flaviviruses^27^ but is higher than that of the PreAm-ZIKV lineage (0.061, 0.047-0.077), likely due to the raised probability of observing slightly deleterious changes in short-term datasets, as observed during previous emerging epidemics^28^.

**Fig. 3.**
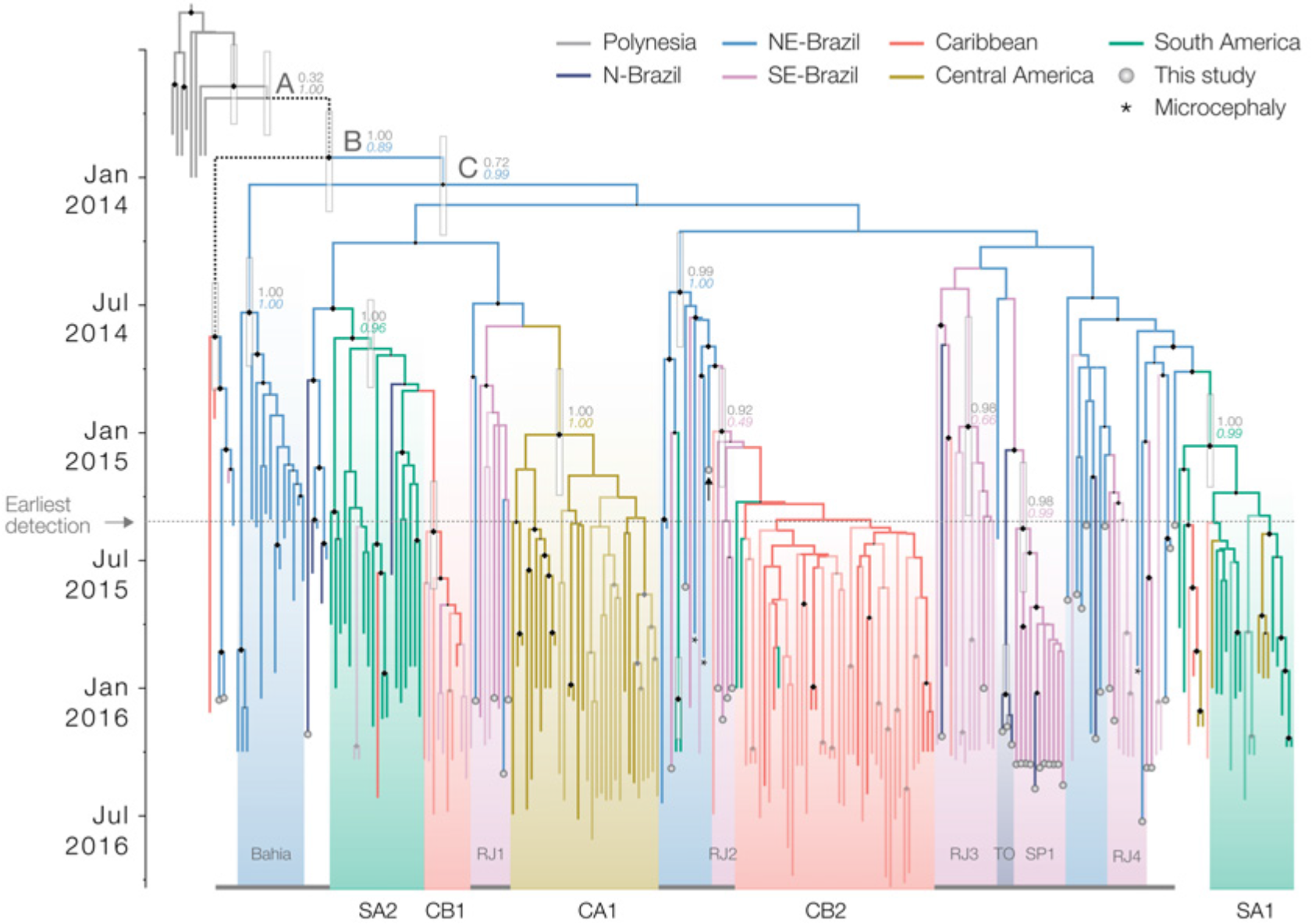
Phylogeography of ZIKV in the Americas. Maximum clade credibility phylogeographic molecular clock phylogeny, estimated from complete and partial Am-ZIKV genomes (see text for details). Terminal branches with grey circles indicate sequences reported in this study. Terminal branches with no circles and reduced opacity represent genomes reported in a companion paper^20^. Thin vertical grey boxes indicate the statistical uncertainty in estimates of the dates of key internal nodes (see Extended Data Table 6). Branch colours indicate the most probable ancestral lineage locations. Size of black diamonds on internal nodes indicate clade posterior probabilities. Coloured numbers show the posterior probabilities of inferred ancestral locations. Asterisks indicate the three available genomes collected from microcephaly cases (Genbank Accession numbers KU497555, KU729217 and KU527068). A black vertical arrow indicates the oldest ZIKV sequence from Brazil. The grey horizontal arrow and horizontal dotted line denotes when ZIKV was first confirmed in the Americas (early May 2015 ^1^). The nodes denoted A and B are equivalent to the nodes named identically in ^4^. Text labels along the bottom of the figure denote clades of viral sequences from regions outside NE Brazil. RJ1 to RJ4 denote clades from Rio de Janeiro state, TO from Tocantins, and SP1 from São Paulo state. Clades from outside Brazil are denoted CB1 and CB2 (Caribbean), SA1 and SA2 (South America excluding Brazil), and CA1 (Central America). The thin grey horizontal lines along the bottom of the figure denote sequences from Brazil.

We used two phylogeographic approaches with different assumptions^29,30^ to reconstruct the spatial origins and spread of ZIKV in Brazil and the Americas. We dated the common ancestor of ZIKV in the Americas (node B, Fig. 3) to Dec 2013 (95% BCIs = Sep 2013-Feb 2014; Extended Data Tables 5 and 6), in line with previous estimates^4,25^. We find evidence that NE Brazil played a central role in the epidemic establishment and dissemination of the Am-ZIKV lineage. Our results suggest that NE Brazil is the most probable location of node B (location posterior support =0.89, Fig. 3). However current data cannot exclude the hypothesis that node B was located in the Caribbean, due the presence of two sequences from Haiti in one of its descendant lineages (Fig. 3 dashed branches). More importantly, most Am-ZIKV sequences descend from a radiation of lineages (node C and its immediate descendants; Fig. 3) that is dated to around Jan 2014 (95% BCIs of node C=Nov 2013-March 2014). Node C is more strongly inferred to have existed in NE Brazil (location posterior support =0.99, Fig. 3) than node B. All 20 replicate analyses performed on sub-sampled data sets place node C in Brazil, 14 of which place node C in NE Brazil (Extended Data Fig. 5). Consequently, we conclude that node C reflects the crucial turning point in the emergence of ZIKV in the Americas. If further data show that node B did indeed exist in Haiti, then it is likely that Haiti acted as an intermediate ‘stepping stone’ for Am-ZIKV’s arrival and establishment in Brazil, from where the virus subsequently spread to other regions. This perspective is consistent with the lower population size of Haiti compared to Brazil (which receives 6 million annual visitors). We infer that node C was present in NE Brazil several months before three notable events that also all occurred in NE Brazil: (i) the retrospective identification of a cluster of suspected but unconfirmed ZIKV cases in Dec 2014^1^, (ii) the oldest ZIKV genome sequence from Brazil, reported here, sampled in Feb 2015, and (iii) confirmed cases of ZIKV transmission in NE Brazil from Mar 2015^31,32^.

**Fig. 4.**
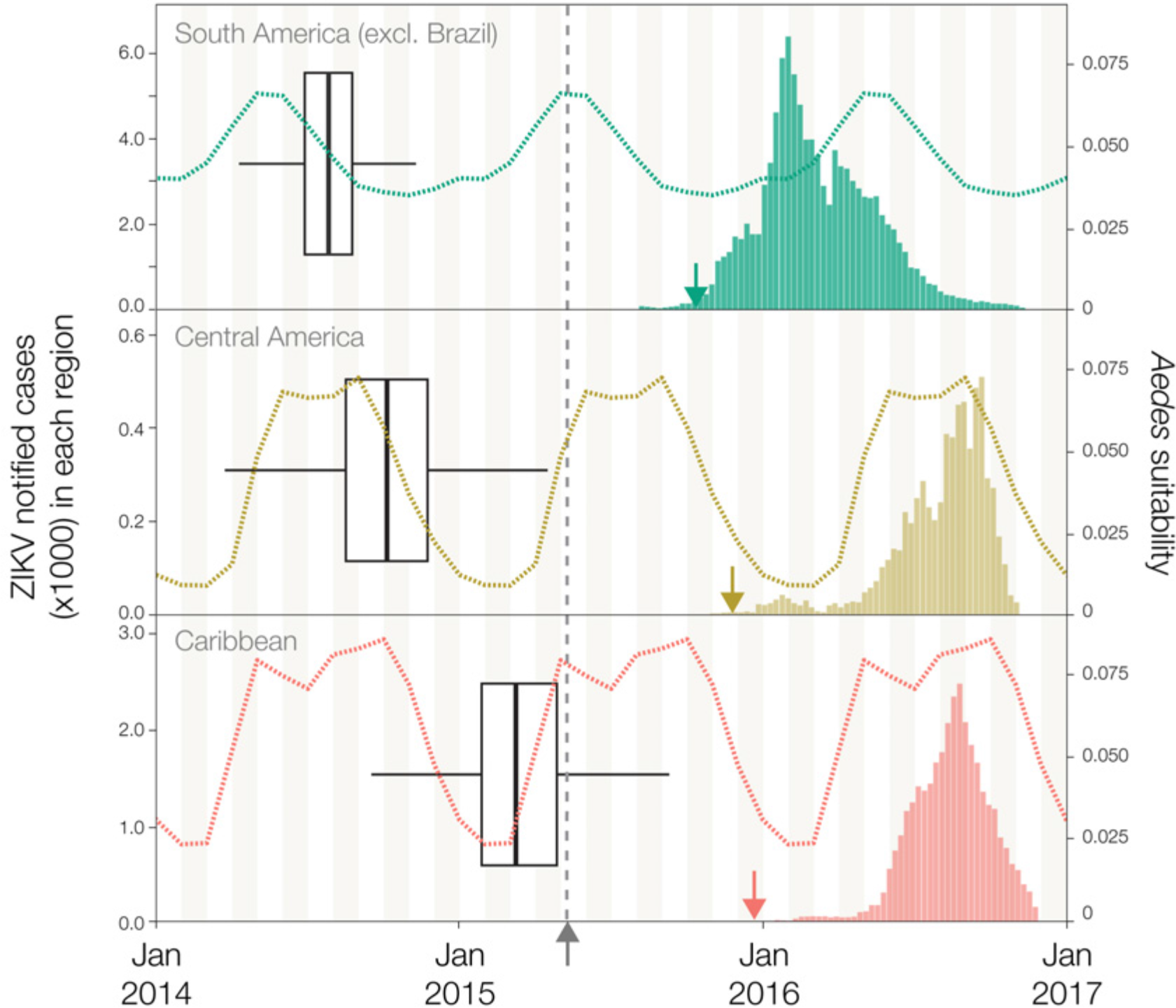
Establishment of Am-ZIKV in the Americas. The earliest inferred dates of lineage export to each non-Brazilian region are represented by box-and-whisker plots. Each refers to the earliest estimated movement between a pair of locations involved in well-supported virus lineage migration. The whiskers and box edges, from left to right, represent the 2.5%, 25%, 75% and 97.5% percentiles of the estimated date of earliest movement. Panel **a** shows the earliest export to South America outside Brazil (SA1 in Fig. 3), **b** shows the export to Central America (CA1), and **c** shows an export to the Caribbean (CB1). In each of **a**-**c**, dashed lines show the estimated climatic vector suitability score for each recipient region, averaged across the countries for which sequence data is available (see **Methods**). In each of **a**-**c**, the bar plots show available notified ZIKV case data from the Pan American Health Organization the countries with the earliest confirmed cases in each non-Brazilian region: Colombia^61^ for South America, Mexico^62^ for Central America, and Puerto Rico^63^ for the Caribbean region. Coloured arrows indicate the earliest date of confirmation of ZIKV autochthonous cases in each non-Brazilian region. The vertical grey dashed line represents the date of ZIKV confirmation in the Americas. The colour scheme is identical to Fig. 3.

Our results further suggest that node C viruses from NE Brazil were important in the continental spread of the epidemic. Within Brazil, we find several instances of virus lineage movement from NE to SE Brazil; most of these events are dated to the second half of 2014, and led to onwards transmission in Rio de Janeiro (RJ1 to RJ4 in Fig. 3) and São Paulo states (SP1 in Fig. 3). We also infer that ZIKV lineages disseminated from NE Brazil to elsewhere in Central America, the Caribbean, and South America. Most Am-ZIKV strains sampled outside Brazil fall into four well-supported phylogenetic groups in Fig 3; three (SA1, SA2/CB1, CA1) are inferred to have been exported from NE Brazil between Jul 2014 and Apr 2015, while the Caribbean clade CB2 appears to originate from SE Brazil between Jan and Mar 2015 (Figs. 3, 4). Each independent viral lineage export occurred during a period of climatic suitability for vector transmission in the recipient location (Fig. 4). For the earliest exports to Central America (CA1) and South America (SA2), there is a 7-11 month gap between the estimated date of exportation and the date of ZIKV detection in the recipient location, suggesting a complete or partial season of undetected transmission. These periods of cryptic transmission are relevant to studies of spatio-temporal trends in reported microcephaly in the Americas, because they help define the appropriate timeframe for baseline (pre-ZIKV) microcephaly in each region.

Combining virus genomic and epidemiological data can generate insights into the patterns and drivers of vector-borne virus transmission. Large-scale surveillance of ZIKV is challenging because (i) many cases may be asymptomatic and (ii) ZIKV co-circulates in some regions with other arthropod-borne viruses that exhibit overlapping symptoms (e.g. dengue, Chikungunya, Mayaro, and Oropouche viruses). A system of continuous and proportional virus sequencing, integrated with surveillance data, could provide timely information to inform effective response and control measures against Zika and other viruses, including the recently re-emerged yellow fever virus^33^.

## Methods

### Sample collection

Between the 1^st^ and 18^th^ June 2016, 1330 samples from cases notified as ZIKV infected were tested for ZIKV infection in the Northeast region of Brazil (NE Brazil). During this period, 4 of the 5 laboratories in the region visited by the ZiBRA project were in the process of implementing molecular diagnostics for ZIKV. The ZiBRA team spent 2-3 days in each state central public health laboratory (LACEN). The samples analysed had been previously collected from patients who had attended a municipal or state public health facility, presenting maculopapular rash and at least two of the following symptoms: fever, conjunctivitis, polyarthralgia, or periarticular edema. The majority of samples were linked to a digital record that collated epidemiological and clinical data: date of sample collection, location of residence, demographic characteristics, and date of onset of clinical symptoms (when available). The ZiBRA project was conducted under the auspices of the *Coordenação Geral de Laboratórios de Saúde Pública* in Brazil (CGLAB), part of the Brazilian Ministry of Health (MoH) in support of the emergency public health response to Zika. Urine and plasma samples from Rio de Janeiro were obtained from patients followed at the Fiocruz Viral Hepatitis Ambulatory (Oswaldo Cruz Institute, Rio de Janeiro, Brazil) following Institutional Review Board approval (IRB142/01) from Oswaldo Cruz Institute. RNA was extracted at the Paul-Ehrlich-Institut and sequenced at the University of Birmingham, UK.

### Nucleic acid isolation and RT-qPCR

Serum, blood and urine samples were obtained from patients 0 to 228 days after first symptoms (Extended Data Table 1). Viral RNA was isolated from 200 ul Zika-suspected samples using either the NucliSENS easyMag system (BioMerieux, Basingstoke, UK) (Ribeirão Preto samples), the ExiPrep Dx Viral RNA Kit (BIONEER, Republic of Korea) (Rio de Janeiro samples) or the QIAamp Viral RNA Mini kit (QIAGEN, Hilden, Germany) (all other samples) according to the manufacturer’s instructions. Ct values were determined for all samples by probe-based RT-qPCR against the prM target (using 5’FAM as the probe reporter dye) as previously described^34^. RT-qPCR assays were performed using the QuantiNova Probe RT-qPCR Kit (20 ul reaction volume; QIAGEN) with amplification in the Rotor-Gene Q (QIAGEN) following the manufacturer’s protocol. Primers/probe were synthesised by Integrated DNA Technologies (Leuven, Belgium). The following reaction conditions were used: reverse transcription (50°C, 10 min), reverse transcriptase inactivation and DNA polymerase activation (95°C, 20 sec), followed by 40 cycles of DNA denaturation (95°C, 10 secs) and annealing-extension (60°C, 40 sec). Positive and negative controls were included in each batch; however, due to the large number of samples tested in a short time it was possible only to run each sample without replication.

### Whole genome sequencing

Sequencing was attempted on all positive samples obtained from NE Brazil regardless of Ct value. All samples collected in Brazil that are reported in this study were sequenced with the Oxford Nanopore MinION. Sequencing statistics can be found in Extended Data Table 2. The protocol employed cDNA synthesis with random primers followed by gene specific multiplex PCR and is presented in detail in Quick et al. ^18^. In brief, extracted RNA was converted to cDNA using the Protoscript II First Strand cDNA synthesis Kit (New England Biolabs, Hitchin, UK) and random hexamer priming. ZIKV genome amplification by multiplex PCR was attempted using the ZikaAsianV1 primer scheme and 40 cycles of PCR using Q5 High-Fidelity DNA polymerase (NEB) as described in Quick et al. ^18^. PCR products were cleaned-up using AmpureXP purification beads (Beckman Coulter, High Wycombe, UK) and quantified using fluorimetry with the Qubit dsDNA High Sensitivity assay on the Qubit 3.0 instrument (Life Technologies). PCR products for samples yielding sufficient material were barcoded and pooled in an equimolar fashion using the Native Barcoding Kit (Oxford Nanopore Technologies, Oxford, UK). Sequencing libraries were generated from the barcoded products using the Genomic DNA Sequencing Kit SQK-MAP007/SQK-LSK208 (Oxford Nanopore Technologies). Sequencing libraries were loaded onto a R9/R9.4 flowcell and data was collected for up to 48 hours but generally less. As described ^18^, consensus genome sequences were produced by alignment of two-direction reads to a Zika virus reference genome (strain H/PF/2013, GenBank Accession number: KJ776791) followed by nanopore signal-level detection of single nucleotide variants. Only positions with ≥20x genome coverage were used to produce consensus alleles. Regions with lower coverage, and those in primer-binding regions were masked with N characters. Validation of our sequencing approach on the MinION platform was undertaken by sequencing with MinION the WHO reference strain of Zika virus that was also sequenced using the Illumina Miseq platform^19^; identical consensus sequences were recovered regardless of the MinION chemistry version employed (R7.3, R9 and R9.4) (Extended Data Fig. 2).

### Collation of genome-wide data sets

Our complete and partial genome sequences were appended to a global data set of all available published ZIKV genome sequences (up until January 2017) using an in-house script that retrieves updated GenBank sequences on a daily basis. In addition to the genomes generated from samples collected in NE Brazil during ZiBRA fieldwork, samples were sent directly to University of São Paulo and elsewhere for sequencing. Thirteen genomes from Ribeirão Preto, São Paulo state (SP; SE-Brazil region) and seven genomes from Tocantins (TO; N-Brazil region) were sequenced at University of São Paulo. Nine genomes from Rio de Janeiro (RJ; SE-Brazil region) were sequenced in Birmingham, UK, and added to our dataset. All these genomes were generated using the same primer scheme as the ZiBRA samples collected in NE Brazil ^18^. In addition to these 45 sequences from Brazil, we further included in analysis 9 genomes from ZIKV strains sampled outside of Brazil in order to contextualise the genetic diversity of Brazilian ZIKV, giving rise to a final data set of 54 sequences. Specifically, we included 5 genomes from samples collected in Colombia and 4 new genomes from Mexico, which were generated using the protocols described in refs. ^35^ and ^22^, respectively.

GenBank sequences belonging to the African genotype of ZIKV were identified using the Arboviral genotyping tool (http://bioafrica2.mrc.ac.za/rega-genotype/typingtool/aedesviruses) and excluded from subsequent analyses, as our focus of study was the Asian genotype of ZIKV and the Am-ZIKV lineage in particular. To assess the robustness of molecular clock dating estimates to the inclusion of older sequences, analyses were performed both with and without the P6-740 strain, the oldest known strain of the ZIKV-Asian genotype (sampled in 1966 in Malaysia). Our final alignment comprised the sequences reported in this study (n=54) plus publicly available ZIKV-Asian genotype sequences, as of 1^st^ March 2017 (*n*=115). We also included in our analysis 85 additional genomes from a companion paper ^20^. The dataset used for analysis therefore included sequences from 254 Zika virus isolates, 241 of which from the Americas. Unpublished but publicly available genomes were included in our analysis only if we had written permission from those who generated the data (see **Acknowledgments**).

### Maximum likelihood analysis and recombination screening

Preliminary maximum likelihood (ML) trees were estimated with ExaMLv3 ^36^ using a per-site rate category model and a gamma distribution of among site rate variation. For the final analyses, ML trees were estimated using PhyML^37^ under a GTR nucleotide substitution model^38^, with a gamma distribution of among site rate variation, as selected by jModeltest.v.2 ^39^. An approximate likelihood ratio test was used to estimate branch support^37^. Final ML trees were estimated with NNI and SPR heuristic tree search algorithms; equilibrium nucleotide frequencies and substitution model parameters were estimated using ML^37^ (see Extended Data Fig. 4).

Recombination may impact evolutionary estimates^40^ and has been shown to be present in the ZIKV-African genotype^41^. In addition to restricting our analysis to the Asian genotype of ZIKV, we employed the 12 recombination detection methods available in RDPv4 ^42^ and the Phi-test approach ^43^ available in SplitsTree ^44^ to further search for evidence of recombination in the ZIKV-Asian lineage. No evidence of recombination was found.

Analysis of the temporal molecular evolutionary signal in our ZIKV alignments was conducted using TempEst^45^. In brief, collection dates in the format yyyy-mm-dd (ISO 8601 standard) were regressed against root-to-tip genetic distances obtained from the ML phylogeny. When precise sampling dates were not available, a precision of 1 month or 1 year in the collection dates was taken into account.

To compare the pairwise genetic diversity of PreAm-ZIKV strains from Asia and the Pacific with Am-ZIKV viruses from the Americas, we used a sliding window approach with 300 nt wide windows and a step size of 50 nt. Sequence gaps were ignored; hence the average pairwise difference per window was obtained by dividing the total pairwise nucleotide differences by the total number of pairwise comparisons.

### Molecular clock phylogenetics and gene-specific d_N_/d_S_ estimation

To estimate Bayesian molecular clock phylogenies, analyses were run in duplicate using BEASTv.1.8.4 ^46^ for 30 million MCMC steps, sampling parameters and trees every 3000 steps. We employed a model selection procedure using both path-sampling and stepping stone models^47^ to estimate the most appropriate model combination for Bayesian phylogenetic analysis. The best fitting model was a HKY codon position-structured SDR06 nucleotide substitution model^48^ with a Bayesian skyline tree prior and a strict molecular clock model (Extended Data Table 5). A non-informative continuous time Markov chain reference prior^49^ on the molecular clock rate was used. Convergence of MCMC chains was checked with Tracer v.1.6. After removal of 10% as burn-in, posterior tree distributions were combined and subsampled to generate an empirical distribution of 1,500 molecular clock trees.

To estimate rates of evolution per gene we partitioned the alignment into 10 genes (3 structural genes C, prM, E, and 7 non-structural genes NS1, NS2A, NS2B, NS3, NS4A, NS4B and NS5) and employed a SDR06 substitution model^48^ and a strict molecular clock model, using an empirical distribution of molecular clock phylogenies. To estimate the ratio of nonsynonymous to synonymous substitutions per site (d_N_/d_S_) for the PreAm-ZIKV and the Am-ZIKV lineages, we used the single likelihood ancestor counting (SLAC) method^50^ implemented in HyPhy^51^. This method was applied to two distinct codon-based alignments and their corresponding ML trees which comprised the PreAm-ZIKV and Am-ZIKV sequences, respectively.

### Phylogeographic analysis

We investigated virus lineage movements using our empirical distribution of phylogenetic trees and the sampling location of each ZIKV sequence. The sampling location of sequences collected from returning travellers was set to the travel destination in the Americas where infection likely occurred. We discretised sequence sampling locations in Brazil into the geographic regions defined in main text. The number of sequences per region available for analysis was 10 for N Brazil, 41 for NE Brazil and 54 for SE Brazil. No viral genetic data was available for the Centre-West (CW) and the South (S) Brazilian regions. We similarly discretised the locations of ZIKV sequences sampled outside of Brazil. These were grouped according to the United Nations M49 coding classification of macro-geographical regions. Our analysis included 53 sequences from the Caribbean, 38 from Central America, 17 from Polynesia, 37 from South America (excluding Brazil), 3 from Southeast Asia and 1 from Micronesia. To account for the possibility of sampling bias arising from a larger number of sequences from particular locations, we repeated all phylogeographic analyses using (i) the full dataset (*n*=254) and (ii) ten jackknife resampled datasets (*n*=74) in which taxa from each location (except for Southeast Asia and Micronesia) were randomly sub-sampled to 10 sequences (the number of sequences available for N-Brazil).

Phylogeographic reconstructions were conducted using two approaches; (i) using the asymmetric^52^ discrete trait evolution models implemented in BEASTv1.8.4^46^ and (ii) using the Bayesian structured coalescent approximation (BASTA)^29^ implemented in BEAST2v.2. The latter has been suggested to be less sensitive to sampling biases^53^. For both approaches, maximum clade credibility trees were summarized from the MCMC samples using TreeAnnotator after discarding 10% as burn-in. The posterior estimates of the location of nodes A, B and C (depicted in Fig. 3) from these two analytical approaches (applied to both the complete and jackknifed data sets) can be found in Extended Data Fig. 5.

For the discrete trait evolution approach, we counted the expected number of transitions among each pair of locations (net migration) using the robust counting approach^54,55^ available in BEASTv1.8.4^46^. We then used those inferred transitions to identify the earliest estimated ZIKV introductions into new regions. These viral lineage movement events were statistically supported (with Bayes factors > 3) using the BSSVS (Bayesian stochastic search variable selection) approach implemented in BEASTv.1.8.4^30^. Box plots for node ages were generated using the ggplot256 package in R software^57^.

### Epidemiological analysis

Weekly suspected ZIKV data per Brazilian region were obtained from the Brazilian Ministry of Health (MoH). Cases were defined as suspected ZIKV infection when patients presented maculopapular rash and at least two of the following symptoms: fever, conjunctivitis, polyarthralgia or periarticular edema. Because notified suspected ZIKV cases are based on symptoms and not molecular diagnosis, it is possible that some notified cases represent other co-circulating viruses with related symptoms, such as dengue and Chikungunya viruses. Further, case reporting may have varied among regions and through time. Data from 2015 came from the pre-existing MoH sentinel surveillance system that comprised 150 reporting units throughout Brazil, which was eventually standardised in Feb 2016 in response to the ZIKV epidemic. We suggest that these limitations should be borne in mind when interpreting the ZIKV notified case data and we consider the R_0_ values estimated here to be approximate. That said, our time series of RT-qPCR+ ZIKV diagnoses from NE Brazil qualitatively match the time series of notified ZIKV cases from the same region (Fig. 1b). To estimate the exponential growth rate of the ZIKV outbreak in Brazil, we fit a simple exponential growth rate model to each stage of the weekly number of suspected ZIKV cases from each region separately:

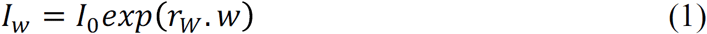
where *I*_*w*_ is the number of cases in week *w*. As described in main text, the Brazilian regions considered here were NE Brazil, N-Brazil, S-Brazil, SE-Brazil, and CW-Brazil. The time period over which exponential growth occurs was determined by plotting the log of *I*_*w*_ and selecting the period of linearity (Extended Data Fig. 6). A linear model was then fitted to this period to estimate the weekly exponential growth rate *r*_*W*_:

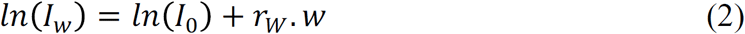
Let *g*(.)be the probability density distribution of the epidemic generation time (i.e. the duration between the time of infection of a case and the mean time of infection of its secondary infections). The following formula can be used to derive the reproduction number R from the exponential growth rate *r* and density *g*(.)^58^.

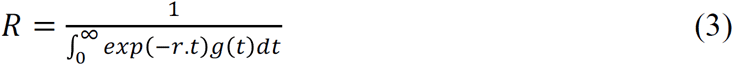

In our baseline analysis, following Ferguson et al.^59^ we assume that the ZIKV generation time is Gamma-distributed with a mean of 20.0 days and a standard deviation (SD) of 7.4 days. In a sensitivity analysis, we also explored scenarios with shorter mean generation times (10.0 and 15.0 days) but unchanged coefficient of variation SD/mean=7.4/20=0.37 (Extended Data Table 3).

### Association between *Aedes aegypti* climatic suitability and ZIKV notified cases

To account for seasonal variation in the geographical distribution of the ZIKV vector *Aedes aegypti* in Brazil we fitted high-resolution maps^60^ to monthly covariate data. Covariate data included time-varying variables, such as temperature-persistence suitability, relative humidity, and precipitation, as well as static covariates such as urban versus rural land use. Maps were produced at a 5km × 5km resolution for each calendar month and then aggregated to the level of the five Brazilian regions used in this study (Extended Data Fig. 7). For consistency, we rescaled monthly suitability values so that the sum of all monthly maps equalled the annual mean map^9^.

We then assessed the correlation between monthly *Aedes aegypti* climatic suitability and the number of weekly ZIKV notified cases in each Brazilian region, to test how well vector suitability explains the variation in the number of ZIKV notified cases. To account for the correlation in each Brazilian region we fit a linear regression model with a lag and two breakpoints. As there may be a lag between trends in suitability and trends in notified cases, we include a temporal term in the model to allow for a shift in the respective curves. Thus for each region, different sets of the constant and linear terms are fitted to different time periods. More formally,

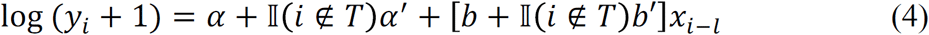
where *y*_*i*_ represents notified cases in a particular region in month *i, x*_*i*_ is the climatic suitability in that region in month *i, l* is the time lag that yields the highest correlation between *y*_*i*_ and *x*_*i*_ and *T* is the set of time indexes in the correlated region.

We then find the values of *T* and *l* that provide the highest adjusted-*R*^2^ by stepwise iterative optimisation. For each value of *T* evaluated, the optimal value of *l* (i.e. that which gives the highest adjusted-*R*^2^ for the model above) is found by the optim function in *R*^57^. Climatic suitability values were only calculated for each month, so to calculate suitability values for any given point in time we interpolated between the monthly values using a linear function. We found no significant effect of residual autocorrelation in our data (Extended Data Fig. 8).

## Data availability

Sequences of the primers and probes used here have been available at http://www.zibraproject.org since the beginning of the project. Genome sequences were made publicly available at http://www.zibraproject.org once generated and confirmed. New Brazilian sequences are available in GenBank under accession numbers KY558989 to KY559033 (see also Extended Data Table 4). The new Colombian and Mexican ZIKV sequences are available under accession numbers KY317936-KY317940 and KY606271 to KY606274, respectively.

## Acknowledgments

We are deeply grateful to Fundação Oswaldo Cruz in Bahia and Pernambuco states, University of São Paulo, and Instituto Evandro Chagas, and to the Brazilian Zika virus surveillance network for their essential contributions to this project. We are grateful to the following researchers for giving us permission to use in our analyses their unpublished genomes available on GenBank: Robert Lanciotti (Centers for Disease Control & Prevention, USA), John Lednicky (University of Florida, USA), Antoine Enfissi (Institut Pasteur de la Guyane), F. Baldanti (Pavia University, Italy), Reed Shabman (ATCC, USA), Brett Picket (J. Craig Venter Institute, USA), Raymond Schinazi (Emory University, USA), Myrna Bonaldo (Instituto Oswaldo Cruz, Rio de Janeiro, Brazil), Michael Gale (University of Washington, USA), Maria Capobianchi and Catilletti Concetta (National Institute for Infectious Diseases “L Spallanzani”), Mariana Leguia (US Naval Medical Research Unity 6, Peru), José Alberto Diaz (InDRE, Mexico), Edgar Sevilla-Reyes (INER, Mexico), Alexander Franz (University of Missouri, USA), Mariano Garcia-Blanco (Duke University, USA) and MJ van Hemert (LUMC, The Netherlands). We thank Pedro Fernando da Costa Vasconcelos, Sueli Guerreiro Rodrigues, Jedson Cardoso, Janaina Vasconcelos, João Vianez Junior (Instituto Evandro Chagas, Brazil), Juliana Gil Melgaço (Fiocruz Rio de Janeiro, Brazil), Johannes Blumel (Paul-Ehrlich-Institut, Langen, Germany), Marcia Cristina Brito Lobato, Liliana Nunes Fava (Tocantins State Department of Health, Brazil) and Constância Ayres (Instituto Aggeu Magalhães, Fundação Oswaldo Cruz (FIOCRUZ), Recife, Pernambuco, Brazil). LCJA thanks QIAGEN for reagents and equipment for the ZiBRA project. MRTN thanks FERPEL for providing consumables. We thank Filipa Campos for advice on Figure 1. We are grateful to the staff of Oxford Nanopore for technical support, with particular thanks to Rosemary Dokos, Zoe McDougall, Simon Cowan, Gordon Sanghera and Oliver Hartwell.

## Funding

This work was supported by the Medical Research Council/Wellcome Trust/Newton Fund Zika Rapid Response Initiative (grant number MC_PC_15100/ ZK/16-078) which also supports JQ’s salary (Grant) and by the generous support of the American people through the United States Agency for International Development Emerging Pandemic Threats Program-2 PREDICT-2 (Cooperative Agreement No. AID-OAA-A-14-00102), which also supports MUGK’s salary. NJL is supported by a Medical Research Council Bioinformatics Fellowship as part of the Cloud Infrastructure for Microbial Bioinformatics (CLIMB) project. NRF is funded by a Sir Henry Dale Fellowship (Wellcome Trust / Royal Society Grant 204311/Z/16/Z). CNPq contributed to the trip expenses (grant no. 457480/2014-9). ACC was supported by FAPESP #2012/03417-7. MRTN is supported by the Brazilian National Council of Scientific and Technological Development (CNPq) grant no. 302584/2015-3. AB and TB were supported by NIH award R35 GM119774. AB is supported by the National Science Foundation Graduate Research Fellowship Program under Grant No. DGE-1256082. TB is a Pew Biomedical Scholar. CYC is partially supported by NIH grant R01 HL105704 and a pathogen discovery award from Abbott Laboratories, Inc. EH is supported by a National Health and Medical Research Council Australia Fellowship (GNT1037231). SCH is supported by Wellcome Trust Grant 102427. This research received funding from the ERC under the European Union’s Seventh Framework Programme (FP7/2007-2013)/ERC, grant agreement numbers 614725-PATHPHYLODYN and 278433-PREDEMICS, and from European Union Horizon 2020 under grant agreements 643476-COMPARE and 734548-ZIKAlliance (to SC). TJ and ETJM and acknowledge funding from IDAMS, DENFREE, DengueTools, and from PPSUS-FACEPE (project Number APQ-0302-4.01/13). RFF received funding from FACEPE, grant number: APQ-0044.2.11/16 and APQ-0055.2.11/16 and from CNPq 439975/2016-6. SAB was supported by the Sicherheit von Blut und Geweben hinsichtlich der Abwesenheit von Zikaviren from the German Ministry of Health.

## Conflicts of Interest

NJL received an honorarium for speaking at an Oxford Nanopore meeting. NJL has ongoing research collaborations with Oxford Nanopore Technologies and has received free-of-charge reagents in support of the ZiBRA project. OGP receives consultancy income from Metabiota Inc, CA, USA. CYC is the director of the UCSF-Abbott Viral Diagnostics and Discovery Center and receives research support from Abbott Laboratories, Inc.

## Supplementary Information

**Extended Data Fig. 1.**
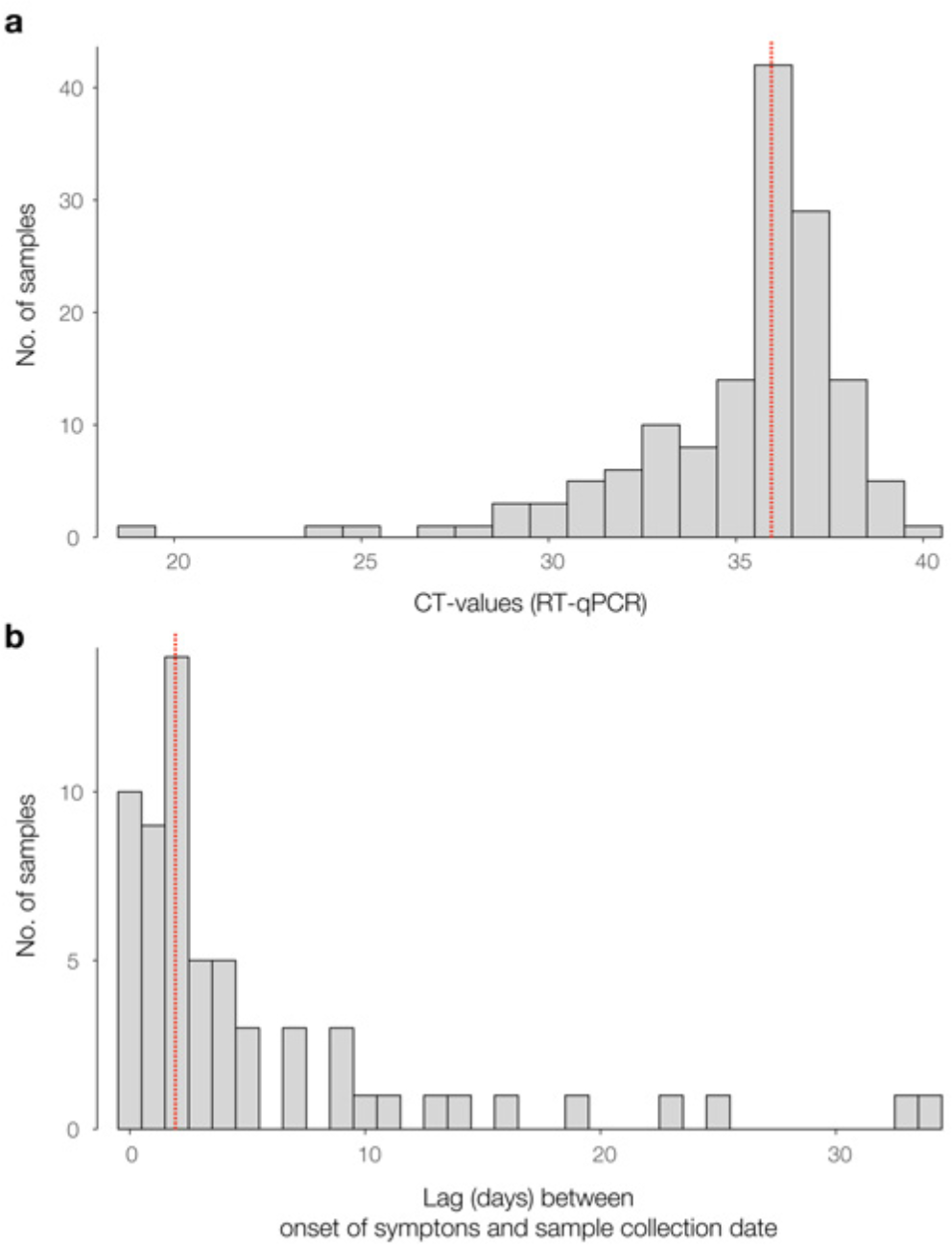
Panel **a** shows the distribution of RT-qPCR+ samples tested during the ZiBRA journey in Brazil (*n*=181 samples; median = 35.96). Panel **b** shows the distribution of the temporal lag between the date of onset of clinical symptoms and the date of sample collection of RT-qPCR+ samples (median = 2 days). Red dashed lines represent the median of the distributions.

**Extended Data Fig. 2.**
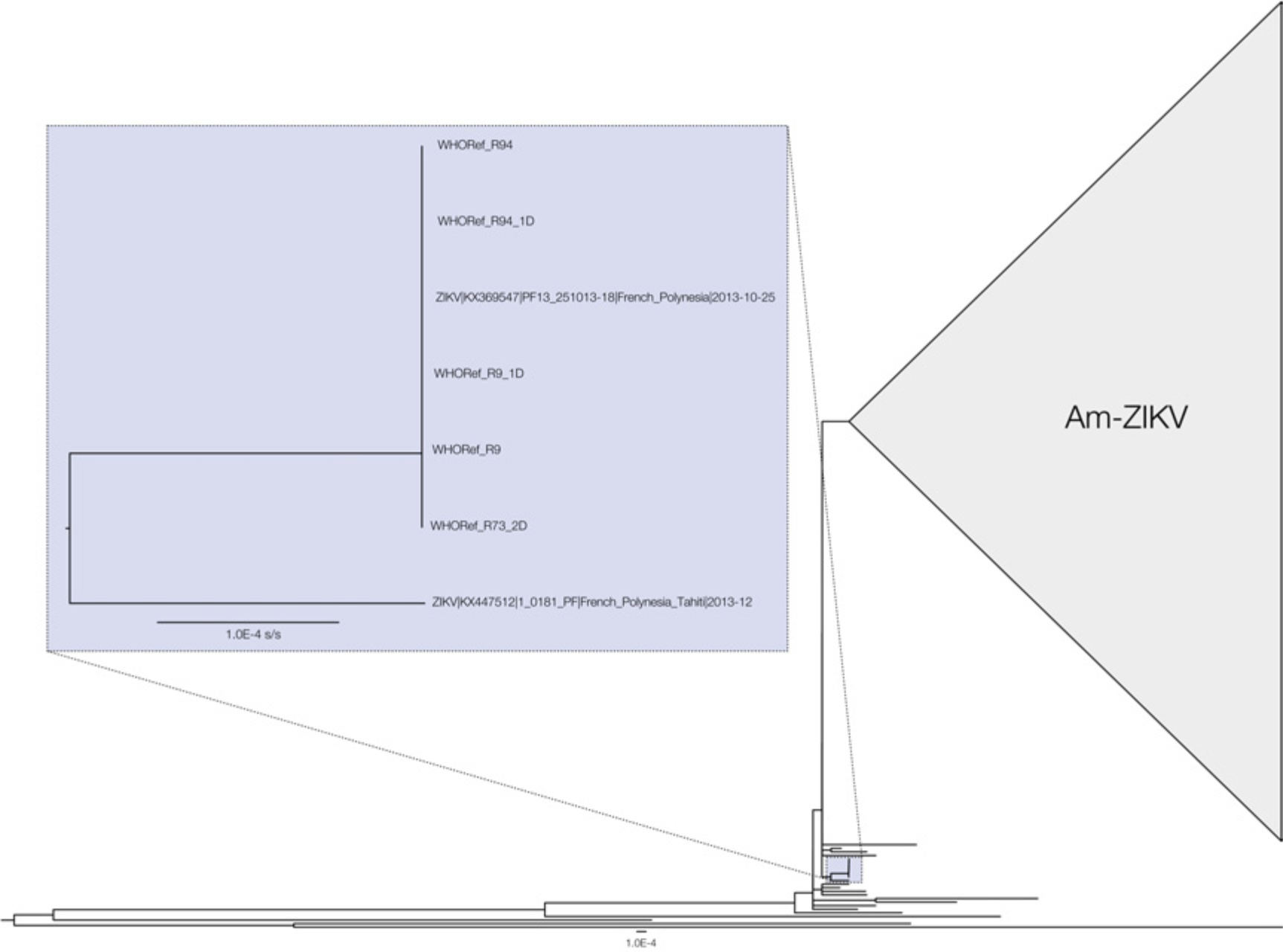
Validation of sequencing approaches. The expanded clade highlighted in blue contains the WHO reference ZIKV sequence^19^ (accession number KX369547) generated using Illumina MiSeq. Sequences generated using MinION chemistries R9.4 2D, R9.4 1D, R9 1D, R9 2D and R7.3 2D are identical and therefore also placed in this clade. The phylogeny was estimated using PhyML^64^. Scale bar represents expected nucleotide substitutions per site (s/s). Am-ZIKV=American Zika virus lineage.

**Extended Data Fig. 3.**
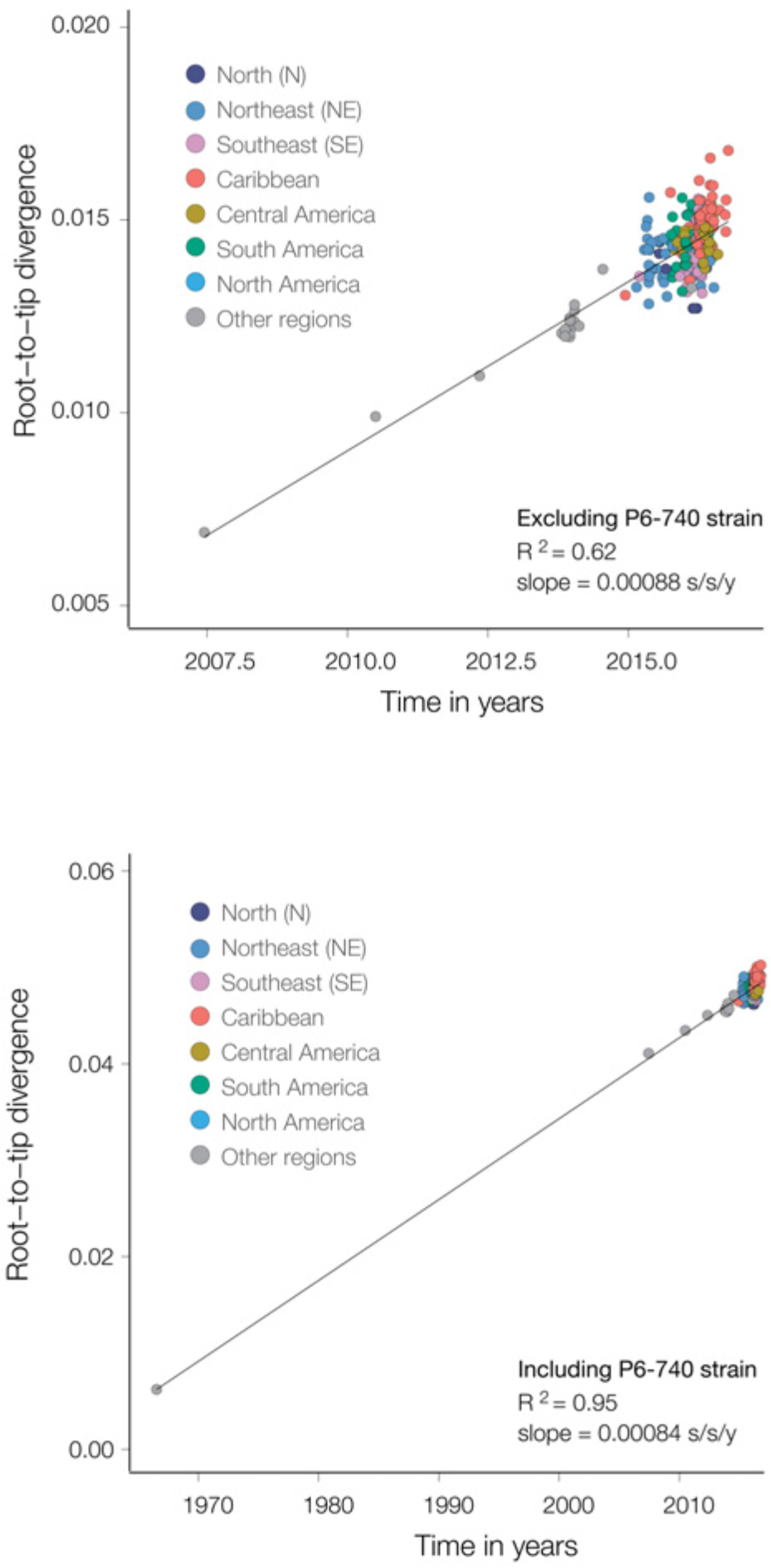
Temporal signal of the ZIKV Asian genotype. The correlation between sampling dates and genetic distances from the tips to the root of a maximum likelihood (ML) tree estimated PhyML^64^ was explored using TempEst^45^. (**a**) Estimates for the dataset used for the phylogenetic analysis in Fig. 3c, and (**b**) estimates for the same dataset including the P6-740 strain sampled in 1966 (accession number HQ234499).

**Extended Data Fig. 4.**
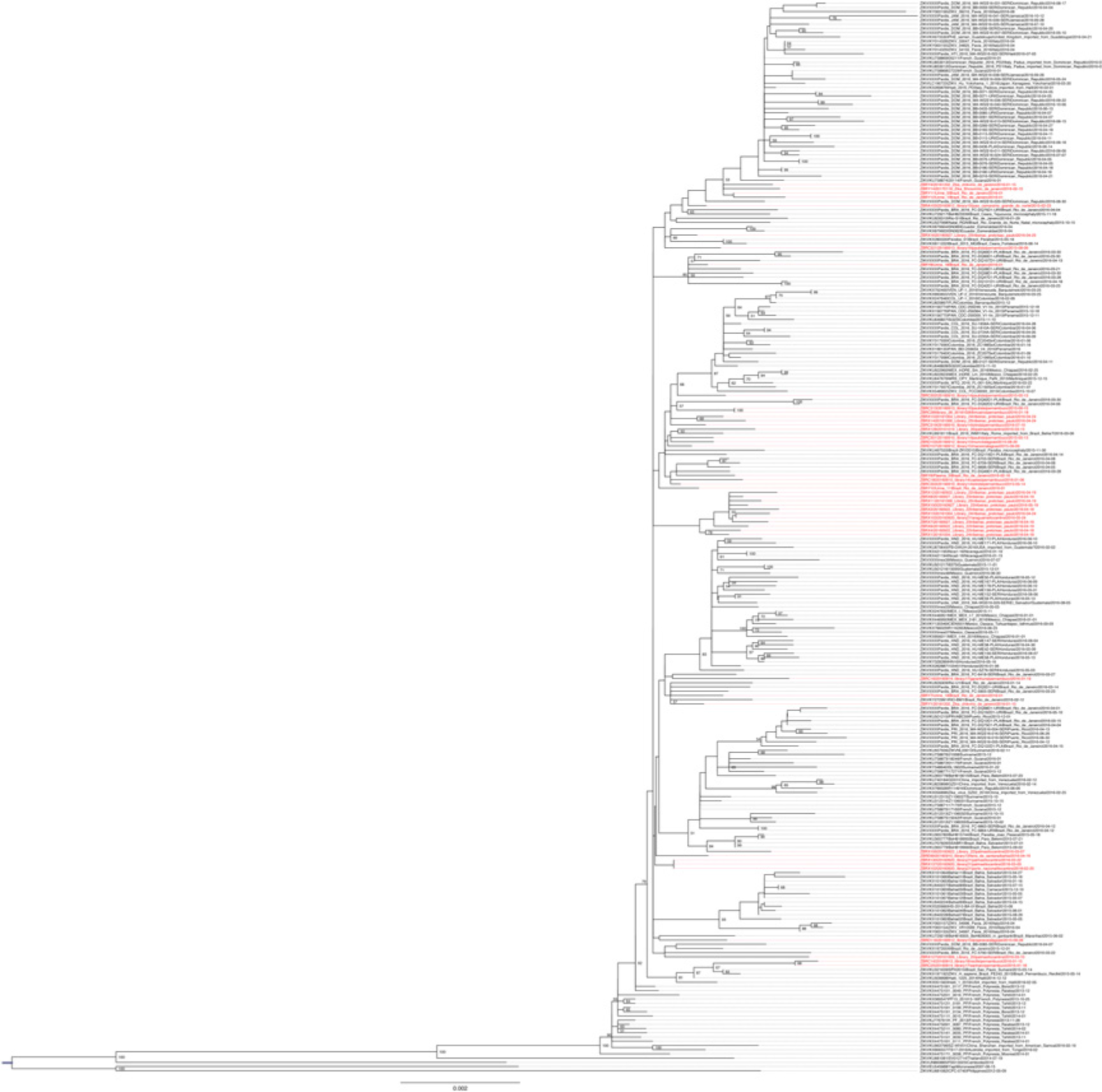
A non-clock maximum likelihood phylogeny of our ZIKV data set. Bootstrap branch support values are shown at each node. The phylogeny was estimated using PhyML^64^. Sequences generated in this study are highlighted in red. Scale bar is shown in units of nucleotide substitutions per site.

**Extended Data Fig. 5.**
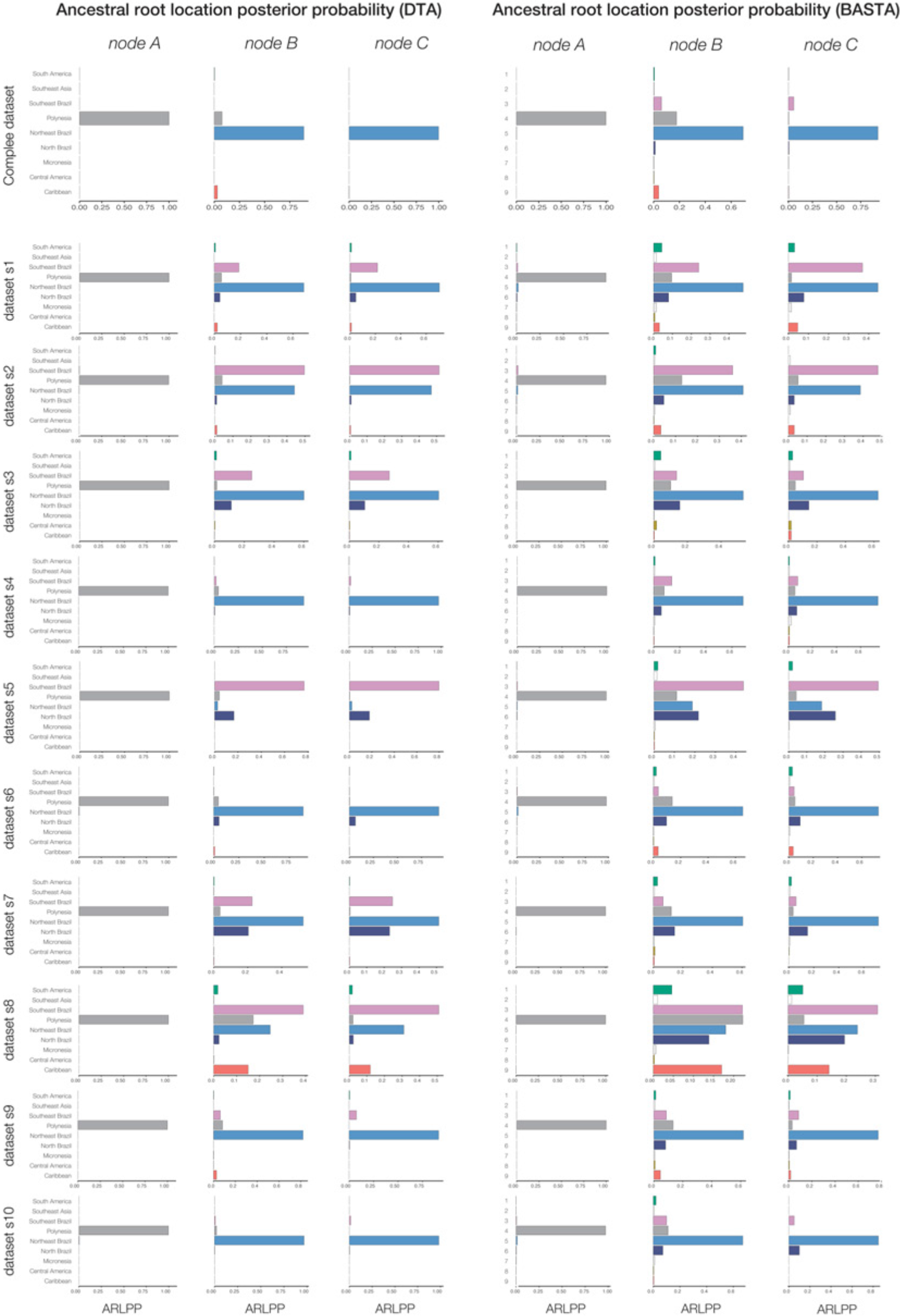
Posterior probabilities of the locations of nodes A, B and C, estimated using the complete dataset (upper panel) and ten replicate subsampled data sets (see **Methods**). DTA=discrete trait analysis method^30^. BASTA=Bayesian structured coalescent approximation method^29^. For each method, we employed an asymmetric model of location exchange to estimate ancestral node locations and to infer patterns of virus spread among regions.

**Extended Data Fig. 6.**
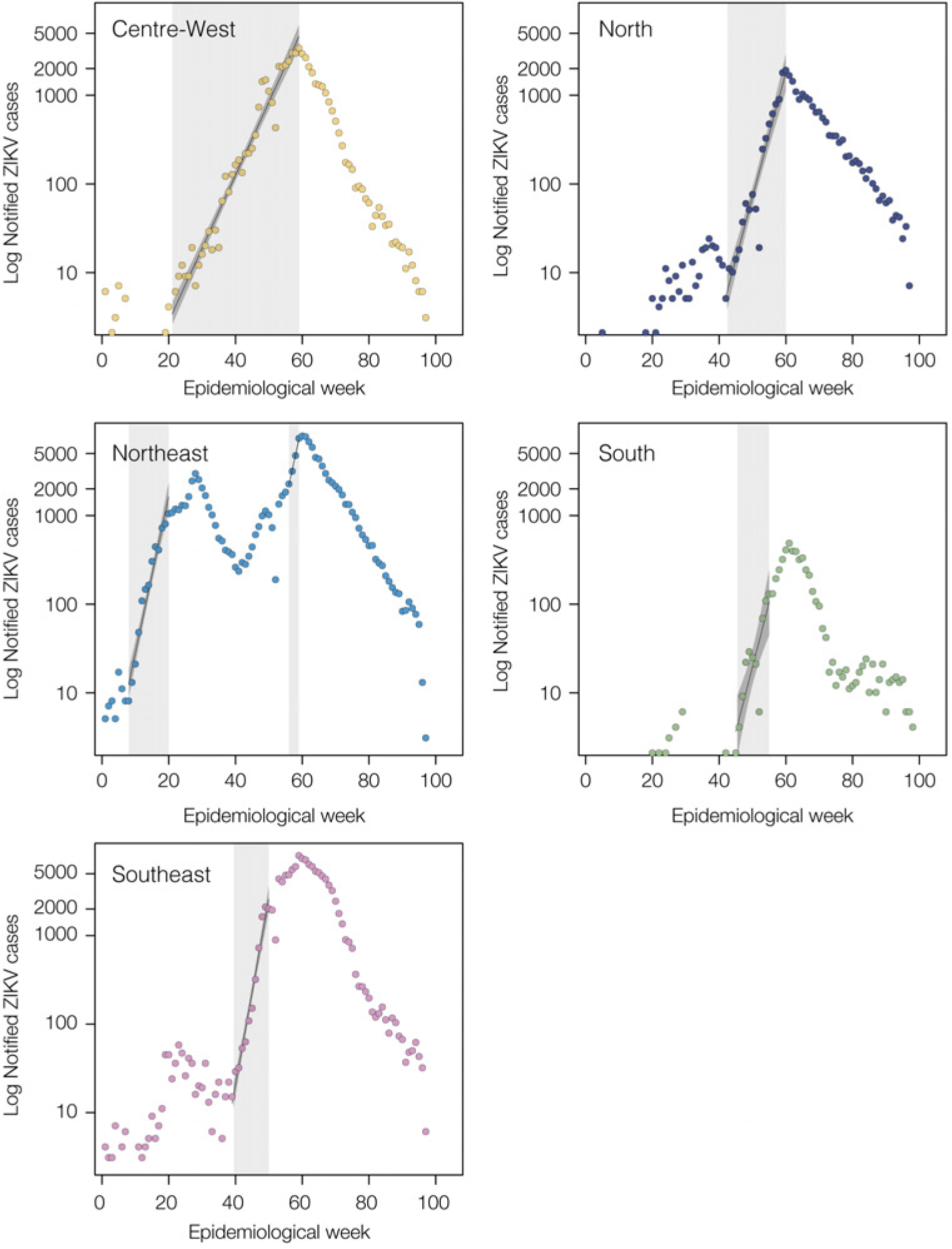
Epidemic growth rates estimated from weekly ZIKV notified cases in Brazil. Time series show the number of ZIKV notified cases in each region of Brazil. Periods from which exponential growth were estimated are highlighted in grey.

**Extended Data Fig. 7.**
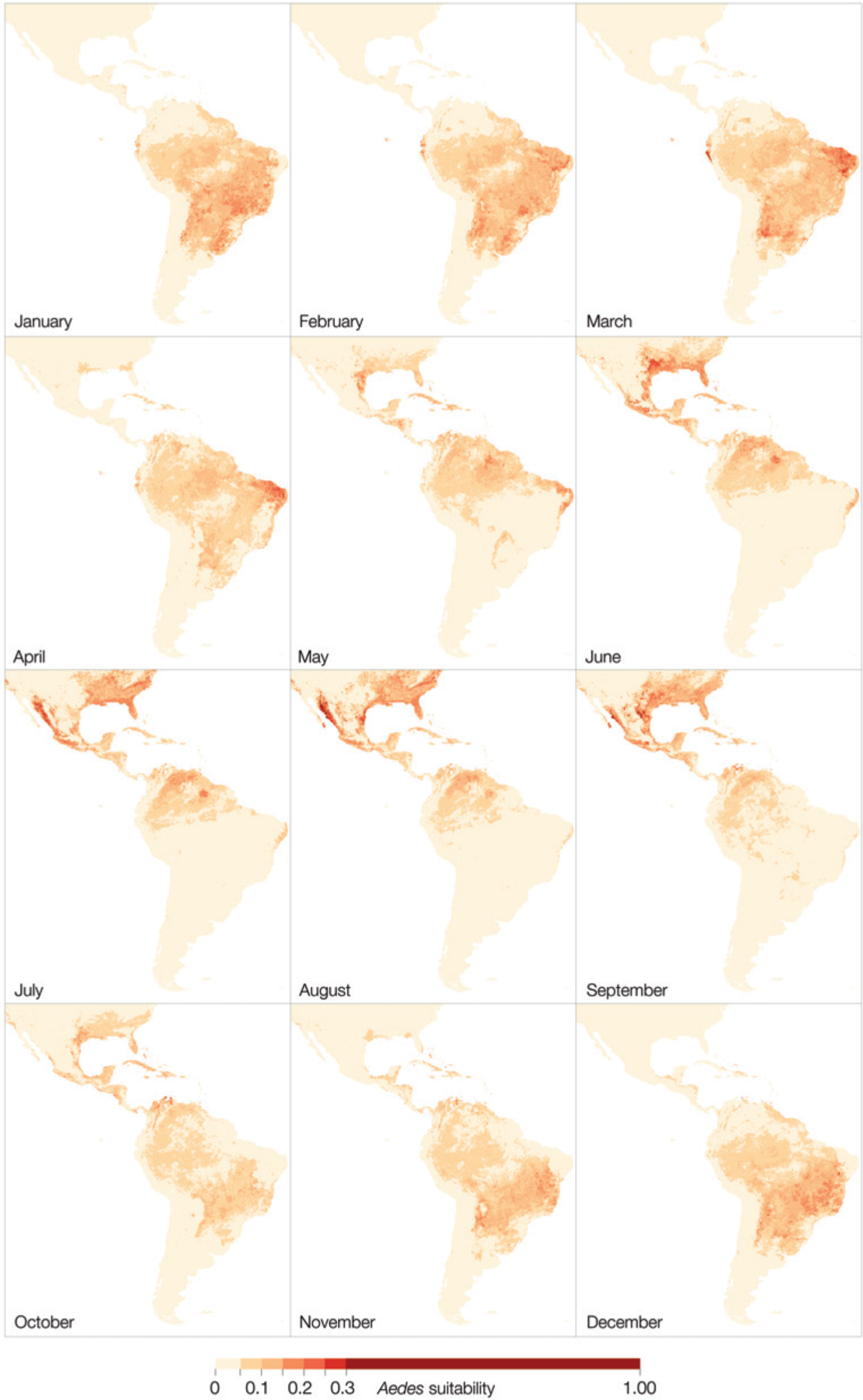
Seasonal suitability for ZIKV transmission in the Americas. These maps were estimated by collating data on *Aedes* mosquitoes, temperature, relative humidity and precipitation, and are the basis of the trends in suitability for different regions shown in main text Figs. 1 and 4. For details, see ^9,60^.

**Extended Data Fig. 8.**
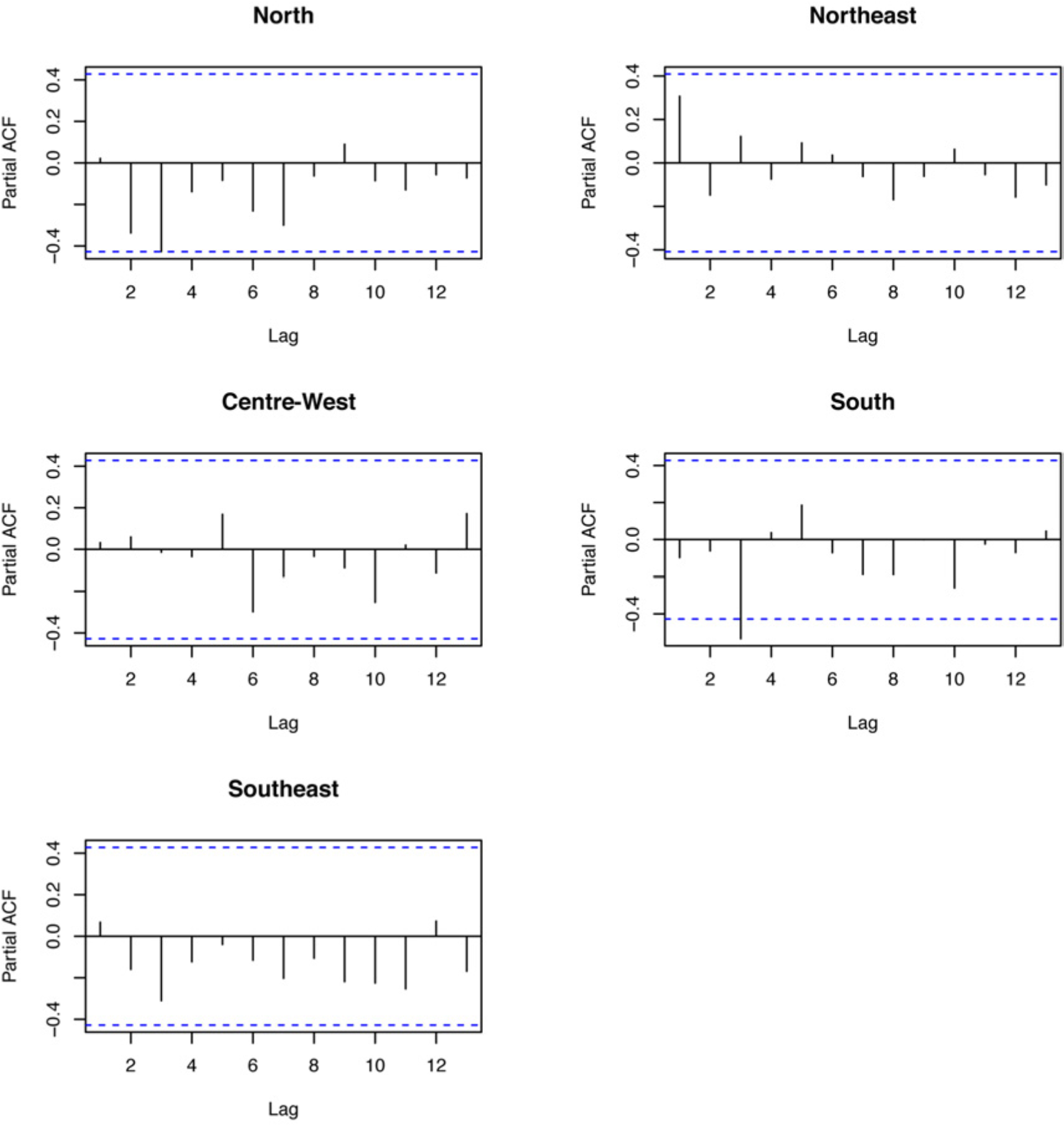
Partial autocorrelation functions for the linear model associating climatic suitability and ZIKV notified cases in each geographic region in Brazil. The residuals for the North, Northeast, Centre-West and Southeast regions show no autocorrelation, while a small amount of autocorrelation cannot be excluded for the South region.

## Extended Data Tables

**Extended Data Table 1.** Summary of the clinical samples tested (*n*=1330, of which 181 were RT-qPCR positive) by the ZiBRA mobile lab in June 2016, NE Brazil. ZIKV notified cases were confirmed using RT-qPCR (see Methods). The collection lag represents the median time interval (in days) between the date of onset of clinical symptoms and the date of sample collection (both dates available for *n*=219) for all samples (including those that subsequently tested RT-qPCR negative). Northeast Brazilian states where samples were tested were RN: Rio Grande do Norte, PB: Paraíba, PE: Pernambuco, AL: Alagoas, BA: Bahia.

| Laboratory,<br>Federal state | No. Positives /<br>Tested (%) | Ct value (mean,<br>min-max) | Collection lag<br>(median, min-max) |
| --- | --- | --- | --- |
| LACEN, RN | 27/335 (8.1%) | 35.9 (18.6-39.1) | 5 (4-16) |
| LACEN, PB | 26/276 (9.4%) | 35.7 (30.7-37.0) | 6 (0-88) |
| FioCruz, PE | 95/315 (30%)<sup>1</sup> | 34.6 (24.1-38.3) | 2.5 (0-33) |
| LACEN, AL | 16/140 (11%) | 34.1 (27.1-40.2) | 2 (0-3) |
| FioCruz, BA | 17/264 (6.4%) | 35.8 (24.7-39.2) | 4 (0-228) |

<sup>1</sup> Includes RT-PCR+ cases from Pernambuco that were generated at Fiocruz Pernambuco.

**Extended Data Table 2.** Parameters of the model measuring the link between climatic vector suitability and notified ZIKV cases in different Brazilian regions. For each region the table provides the estimated correlated time period (*T*), P-value of the linear term of suitability in *T*, adjusted-*R*^2^ of the model, and time lag (*l*).

|  | **North** | **Northeast** | **Centre-West** | **South** | **Southeast** |
| --- | --- | --- | --- | --- | --- |
| **Correlated time period** | 12/2015 to 10/2016 | 7/2015 to 10/2016 | 9/2015 to 8/2016 | 6/2015 to 05/2016 | 11/2015 to 9/2016 |
| **P-value** | <0.0001 | 0.00013 | <0.0001 | <0.0001 | <0.0001 |
| **Adjusted-$R^2$** | 0.929 | 0.8448 | 0.987 | 0.9543 | 0.953 |
| **Time lag (months)** | 1.27 | 0 | 1.12 | 1.19 | 1.33 |

**Extended Data Table 3.** For each region, estimates of the basic reproductive number (*R*) of ZIKV are shown for several values of generation time (*g*) parameter, together with the corresponding estimates of exponential growth rate (*r*) (per day) obtained from notified ZIKV case counts (see also Extended Data Fig. 7). CW: Centre-West, N: North, NE: Northeast (1st: epidemic wave, in 2015; 2nd: epidemic wave, in 2016), SE: Southeast, S: South. CI: 95% confidence interval.

| **Region** | **$R$ (mean, CI),**<br>**$g=20$ days** | **$R$ (mean, CI),**<br>**$g=15$ days** | **$R$ (mean, CI),**<br>**$g=10$ days** | **Growth rate ($r$,**<br>**CI)** |
| --- | --- | --- | --- | --- |
| **CW** | 1.71<br>(1.65-1.78) | 1.46<br>(1.20-1.77) | 1.29<br>(1.13-1.46) | 0.027<br>(0.02-0.03) |
| **N** | 2.48<br>(2.19-2.81) | 1.98<br>(1.80-2.18) | 1.58<br>(1.48-1.69) | 0.046<br>(0.04-0.05) |
| **NE, 1<sup>st</sup>** | 3.12<br>(2.69-3.60) | 2.36<br>(2.11-2.63) | 1.78<br>(1.65-1.91) | 0.06<br>(0.05-0.07) |
| **NE, 2<sup>nd</sup>** | 3.03<br>(2.74-3.36) | 2.31<br>(2.14-2.49) | 1.75<br>(1.66-1.84) | 0.06<br>(0.05-0.06) |
| **SE** | 3.85<br>(3.35-4.42) | 2.77<br>(2.49-3.07) | 1.98<br>(1.84-2.12) | 0.07<br>(0.06-0.076) |
| **S** | 2.57<br>(1.72-3.82) | 2.04<br>(1.50-2.75) | 1.61<br>(1.31-1.97) | 0.05<br>(0.04-0.07) |

**Extended Data Table 4.** Sequencing statistics. Accession numbers, sample IDs, sequencing coverage, RT-qPCR values and epidemiological information for the samples from Brazil generated in this study.

| Accession Number | Sample ID | Aligned Reads | Consensus nucleotide bases (% of reference) | RT-qPCR Ct | Collection Date | Municipality | State |
| --- | --- | --- | --- | --- | --- | --- | --- |
| KY558989 | ZBRA105 | 58128 | 9846 (92) | 29.5 | 2015-02-23 | João Câmara | RN |
| KY558990 | ZBRC14 | 19111 | 8612 (81) | 32.81 | 2016-01-15 | Recife | PE |
| KY558991 | ZBRC16 | 9161 | 7178 (67) | 34.94 | 2016-01-19 | Garanhuns | PE |
| KY558992 | ZBRC18 | 7183 | 7459 (70) | 35.14 | 2016-01-06 | Caetes | PE |
| KY558993 | ZBRC25 | 20533 | 5688 (53) | 35.89 | 2016-01-18 | Sanharó | PE |
| KY558994 | ZBRC28 | 7905 | 8987 (84) | 36.02 | 2016-01-18 | Limoeiro | PE |
| KY558995 | ZBRC301 | 20826 | 9843 (92) | 31.99 | 2015-05-13 | Paulista | PE |
| KY558996 | ZBRC302 | 26331 | 10007 (94) | 30.78 | 2015-05-13 | Paulista | PE |
| KY558997 | ZBRC303 | 12575 | 5873 (55) | 32.81 | 2015-05-14 | Olinda | PE |
| KY558998 | ZBRC313 | 16530 | 9478 (89) | 30.77 | 2015-06-15 | Paulista | PE |
| KY558999 | ZBRC319 | 17316 | 10565 (99) | 24.07 | 2016-07-10 | Olinda | PE |
| KY559000 | ZBRC321 | 11434 | 8647 (81) | 30.62 | 2015-08-09 | Paulista | PE |
| KY559001 | ZBRD103 | 13192 | 8380 (78) | 29.09 | 2015-08-20 | Murici | AL |
| KY559002 | ZBRD107 | 77118 | 7415 (69) | 30.31 | 2015-09-09 | Maceió | AL |
| KY559003 | ZBRD116 | 21211 | 9785 (92) | 27.13 | 2015-08-28 | Arapiraca | AL |
| KY559004 | ZBRE69 | 2313 | 6866 (64) | 24.72 | 2016-04-16 | Feira de Santana | BA |
| KY559005 | ZBRX1 | 21267 | 10559 (99) | 25 | 2016-04-18 | Ribeirão Preto | SP |
| KY559006 | ZBRX2 | 24105 | 9961 (93) | 32 | 2016-04-18 | Ribeirão Preto | SP |
| KY559007 | ZBRX4 | 14722 | 10563 (99) | 26 | 2016-04-18 | Ribeirão Preto | SP |
| KY559008 | ZBRX6 | 12516 | 6893 (64) | 33 | 2016-04-19 | Ribeirão Preto | SP |
| KY559009 | ZBRX7 | 10981 | 8563 (80) | 33 | 2016-04-19 | Ribeirão Preto | SP |
| KY559010 | ZBRX8 | 7445 | 8702 (81) | 33 | 2016-04-19 | Ribeirão Preto | SP |
| KY559011 | ZBRX11 | 21214 | 9379 (88) | 31 | 2016-04-19 | Ribeirão Preto | SP |
| KY559012 | ZBRX12 | 19838 | 10305 (97) | 31 | 2016-04-19 | Ribeirão Preto | SP |
| KY559013 | ZBRX13 | 11809 | 10564 (99) | 21 | 2016-04-24 | Ribeirão Preto | SP |
| KY559014 | ZBRX14 | 5873 | 7469 (70) | 33 | 2016-04-24 | Ribeirão Preto | SP |
| KY559015 | ZBRX15 | 20190 | 10563 (99) | 27 | 2016-04-24 | Ribeirão Preto | SP |
| KY559016 | ZBRX16 | 9698 | 9027 (85) | 32 | 2016-04-25 | Ribeirão Preto | SP |
| KY559017 | ZBRX100 | 5976 | 9609 (90) | 28.5 | 2016-05-19 | Ribeirão Preto | SP |
| KY559018 | ZBRX102 | 13990 | 9508 (89) | 33.91 | 2016-02-25 | Porto Nacional | TO |
| KY559019 | ZBRX103 | 17635 | 9514 (89) | 36.76 | 2016-05-24 | Araguaina | TO |
| KY559020 | ZBRX106 | 29877 | 8458 (79) | 32.36 | 2016-03-07 | Palmas | TO |
| KY559021 | ZBRX127 | 18914 | 10066 (94) | 29.6 | 2016-03-10 | Palmas | TO |
| KY559022 | ZBRX128 | 18480 | 8650 (81) | 28.79 | 2016-03-13 | Palmas | TO |
| KY559023 | ZBRX130 | 16667 | 9914 (93) | 29.06 | 2016-03-22 | Palmas | TO |
| KY559024 | ZBRX137 | 15895 | 9767 (91) | 34.83 | 2016-03-03 | Palmas | TO |
| KY559025 | ZBRY1 | 41036 | 8941 (84) † | 33.53 | 2016-01 | Rio de Janeiro | RJ |
| KY559026 | ZBRY4 | 27865 | 8433 (79) † | 34.21 | 2016-01 | Rio de Janeiro | RJ |
| KY559027 | ZBRY6 | 11779 | 10300 (97) † | 22.66 | 2016-01 | Rio de Janeiro | RJ |
| KY559028 | ZBRY12 | 4980 | 3061 (28) † | 33.66 | 2016-01 | Rio de Janeiro | RJ |
| KY559029 | ZBRY11 | 18530 | 5873 (55) † | 31.11 | 2016-01 | Rio de Janeiro | RJ |
| KY559030 | ZBRY10 | 14067 | 5712 (53) † | 30.84 | 2016-01 | Rio de Janeiro | RJ |
| KY559031 | ZBRY8 | 5708 | 9184 (86) † | 30.96 | 2016-01 | Rio de Janeiro | RJ |
| KY559032 | ZBRY7 | 7749 | 9018 (84) † | 28.07 | 2016-01 | Rio de Janeiro | RJ |
| KY559033 | ZBRY14 | 8040 | 5389 (50) † | 34.2 | 2016-02-15 | Rio de Janeiro | RJ |

† Alignments were performed against version 2 (KJ776791.2) of the genome reference for these sequences; all others used version 1 (KJ776791.1).

**Extended Data Table 5.** Log-marginal likelihood estimates using the path-sampling (PS) and Stepping-Stone (SS) model selection approaches^47,65^. The overall ranking of the models is shown in parentheses for each estimator and the best-fitting combination is underscored. Two molecular clock models were tested here. SC: Strict clock model, UCLD: uncorrelated relaxed clock with lognormal distribution^66^.

| **Clock** | **Coalescent** | **PS** | **SS** |
| --- | --- | --- | --- |
| <u>SC</u> | <u>Skyline</u> | <u>-32141.784 (1)</u> | <u>-32163.360 (1)</u> |
| SC | Exponential | -32239.562 (2) | -32262.003 (2) |
| SC | Constant | -32240.979 (3) | -32266.426 (3) |
| UCLD | Skyline | -35228.978 (4) | -35253.793 (4) |
| UCLD | Exponential | -35294.060 (5) | -35323.312 (5) |
| UCLD | Constant | -35346.424 (6) | -35376.240 (6) |

**Extended Data Table 6.**
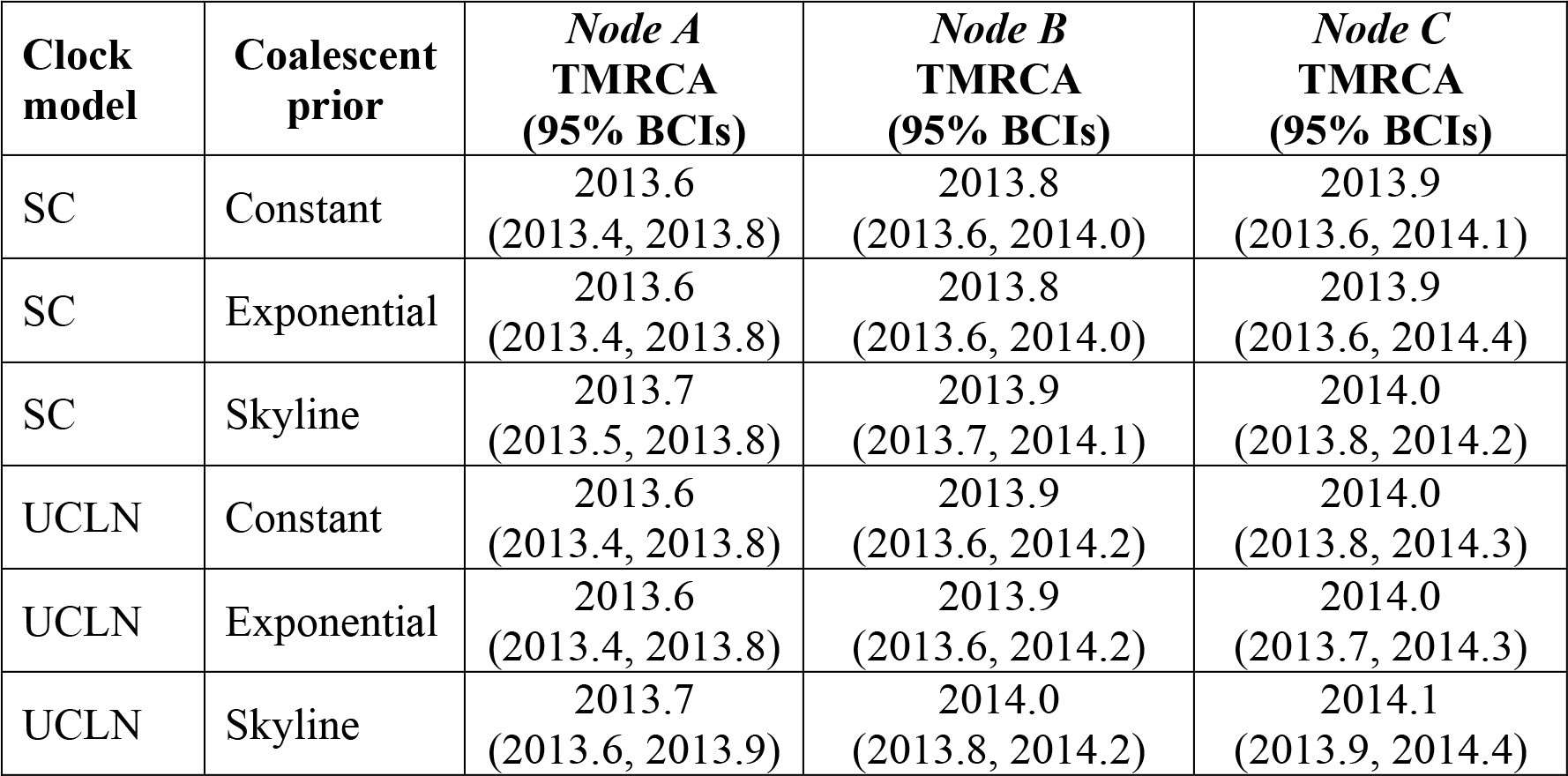
Estimated dates of nodes A, B and C (Fig. 3) under various different molecular clock and coalescent model combinations. TMRCA: time of the most recent common ancestor, BCI: Bayesian credible interval, SC: strict molecular clock model, UCLN: uncorrelated clock with lognormal distribution.

